# Modeling Down syndrome neurodevelopment with isogenic cerebral organoids

**DOI:** 10.1101/2022.05.25.493459

**Authors:** Jan T. Czerminski, Oliver D. King, Jeanne B. Lawrence

## Abstract

As a model of early fetal brain development in Down syndrome, this study examines cortical organoids generated from isogenic trisomic and disomic iPSC lines. Initially pools of organoids from a trisomic versus disomic line found broad transcriptomic differences and modest differences in cell-type representation, suggesting a potential neurodevelopmental phenotype due to Trisomy 21. To better control for multiple sources of variation, we undertook a very robust study of ~1,200 organoids, using an expanded panel of six isogenic subclones (three disomic and three trisomic). The power of the experimental design was indicated by exceptionally strong detection of the ~1.5-fold difference in most chr21 genes. Despite some variability in secreted Aβ-40 levels between “identical” cell lines, this Alzheimer-related phenotype was detected as clearly correlated with Trisomy 21. However, the many statistically significant non-chr21 DEGs found in the small experiment fell away in the expanded study design, such that just three non-chr21 DEGs correlated to T21 status. Similarly, differences in cell-type representation of organoids varied somewhat between the six isogenic lines, but did not correlate with T21 status. Overall, our results indicate that even when organoid and batch variability are better controlled, common, subtle differences between isogenic cell lines (even subclones) may obscure, or be confused with, differences due to Trisomy 21. Interestingly, the neurodegenerative increase in Aβ due to T21 was strong enough to be evident in “fetal” organoids. In contrast, any neurodevelopmental phenotype that may be present in the ~2^nd^ trimester of DS brain development may be more subtle, and within the range of variability in neurodifferentiation potential (unrelated to Trisomy 21) of our isogenic iPSC lines. The potential significance of two non-Chr21 DEGs that results suggest correlate with T21 is discussed.

## Introduction

Cognitive disability is a universal feature of DS, and while the genetic basis of DS is clear, the direct molecular and cellular causes for this phenotype are not well understood. Many studies have raised important hypotheses for specific cell phenotypes and mechanisms, but various findings have not been consistently found, and in some cases are conflicting. For example, several studies have suggested that interneuron number may be decreased in DS patients and human cell models (Ross et al.,1984; Bhattacharyya et al.,2009; Huo et al.,2018), in contrast to other reports in trisomic mice and human organoids that interneuron numbers are increased (Chakrabarti et al.,2010; Das et al.,2013; Xu et al.,2019).

In recent years, with the advent of whole genome sequencing approaches, studies have begun to examine differences in the transcriptomes of DS versus euploid samples, but at this early stage there are few consistent conclusions. Studies do invariably agree that many chr21 genes are upregulated in DS tissues and cells, although the number and identity of these genes is often inconsistent. One meta-analysis of 45 transcriptome studies found only 77 chr21 genes to be consistently upregulated in DS samples (Vilardell et al.,2011) and a more recent meta-analysis of 67 different studies including mouse and human datasets found only 67 “consistently upregulated” genes on chr21 (De Toma et al.,2021). If there is no feedback regulation of a specific gene, then trisomy for a given expressed gene would be expected to result in a 1.5-fold increase in mRNA level, however sensitivity to detect this relatively modest change in each Chr21 gene can be reduced by numerous sources of variation between samples (e.g. genetic background, cell-type proportions in sample, pathological state, age, sex, etc.) Such variables between samples will also broadly influence expression levels of non-chr21 genes which will be particularly dependent on factors that impact cell/tissue state. Thus it remains a challenge to identify consistent changes that are due to trisomy 21.

Most recently, several studies have reported that trisomy 21 causes broad transcriptome-wide changes, with some studies reporting global genomic upregulation genes (Mowery et al.,2018) or the presence of domains of up- and down-regulation across the genome (Letourneau et al.,2014). However, the latter phenomenon was more recently called into question (Do et al.,2015), was not seen in other studies (Gonzales et al.,2018; Moon and Lawrence,2022), or found in both normal and trisomic samples (Ahlfors et al.,2019). Studies in adult DS brain tissue have found hundreds of non-Chr21 genes differentially expressed under stringent statistical cutoffs (Lockstone et al.,2007). However, such differences will reflect differences in cell type proportions in tissue samples, age, or pathological states. For example, some evidence from post-mortem DS brain samples, and human cellular models, find increased numbers of astroglia (Mito and Becker,1993; Lu et al.,2011; Zdaniuk et al.,2011; Briggs et al.,2013; Chen et al.,2014). Such differences in cell-type proportions or tissue status alone could account for broad transcriptome changes in brain samples, complicating identification of specific pathways directly perturbed by trisomy 21.

Numerous molecular pathways and specific Chr21 genes (for example, DYRK1A, RCAN1, Oligo2 or 1, DSCAM, SOD1, Pericentrin) are hypothesized to be central, however consensus has not been reached. Modestly smaller fetal brain sizes have been reported in DS fetuses; however, DS infants often have developmental milestones closer to normal, hence it is important to better understand when in human pre-natal and/or post-natal periods the cognitive deficits arise. This is key to therapeutic strategies, and has been studied in DS mouse models (Bartesaghi et al.,2015; Ruparelia et al.,2012). Recently, new non-invasive *in utero* imaging technologies make studies of human brain development more feasible; for instance, a recent study of DS fetuses by MRI (Patkee et al.,2020) found reduced cerebellar volume in late second trimester compared to controls. While expanding such studies will be important, methods are needed to investigate and better define the cellular neurodevelopmental changes, including cell-types, functions and molecular pathways impacted, information essential for the development of effective therapeutic targets and strategies.

Recently, methods to generate cerebral organoids from human pluripotent stem cells have emerged as a new model system for early human neurodevelopment, modeling brain tissues (Di Lullo and Kriegstein,2017). Organoid systems model development of more complex tissues with a variety of cell types, and over a longer time frame, and protocols for specific brain regions continue to be developed, in what is a promising but young and rapidly evolving research approach.

This study began with the goal of using organoid technology to model the impact of trisomy 21 on human fetal neurodevelopment. While our findings provide new information regarding the extent of early neurodevelopmental changes due to trisomy 21, results and insights from this work are also instructive for the use and interpretation of neurodevelopmental modeling using stem cells. As our efforts that began several years ago evolved, we progressively developed improved experimental design strategies, which we believe are informative more broadly for organoid and stem cell technology to study DS and potentially other neurodevelopmental conditions.

Our lab’s recent DS studies relied on an inducible system to examine the direct effects of “silencing trisomy” in a given trisomic cell population (Chiang et al.,2018; Czerminski and Lawrence,2020; Moon and Lawrence,2022), which avoids comparisons between different isogenic cell lines. Initially when this study began, we had difficulty using the XIST-inducible system in organoids, therefore we identified an all-isogenic panel of trisomic and disomic cell lines for comparison. Results begin with summarizing our test of three organoid generation protocols and choice to utilize a directed forebrain spheroid method for the rest of the study. Having found cytological markers for cell types etc. difficult to quantify in organoids, we conducted in-depth transcriptomics to identify differentially expressed genes and deconvolve cell type representation differences in three trisomic and three isogenic disomic cell lines.

In addition to neurodevelopmental deficits, trisomy 21 causes neurodegeneration as a form of early-onset Alzheimer’s Disease (EOAD), with DS individuals developing amyloid plaques as early as adolescence, and ~80% show clinical dementia by ~60-65 years (Wiseman et al.,2015). While AD-related neuropathologies are not examined in detail, we examined whether an increase in Aβ levels was detected in our experiments, as a comparison to neurodevelopmental related changes, and potential indicator for the sensitivity of experimental design. The APP gene is clearly a driver of AD in DS, since non-trisomic individuals with APP duplication have fully penetrant EOAD (Hithersay et al. 2019; Wiseman et al. 2015). Increased production of Aβ by gamma-secretase (g-sec) cleavage of APP are markers of AD cell pathology prior to dementia (Lehmann et al.,2018; Fortea et al.,2020), hence we briefly examined Aβ secretion in the fetal-stage organoids.

Some methodological points demonstrated by this work have value for the field of disease modeling with human iPSCs more generally, whereas other specific results have significant implications for the fundamental biology of trisomy 21.

## Materials and Methods

### iPSC culture

The isogenic cell lines used here were generated and characterized as described in our prior study (Jiang et al.,2013), and expanded to identify six all isogenic subclones, derived from the same DS iPSC parental line (DS1-iPS4) (Park et al.,2008). In characterizing ~100 subclones for the prior study focused on creating XIST transgenic lines, we identified many subclones that were not transgenic for XIST (but carried the tet-puromycin selection gene). Some subclones were shown to be euploid by spontaneous loss of one chromosome 21, with Chr21 transcriptome levels equivalent to non-isogenic normal control cells (Jiang et al.,2013). Several such trisomic and disomic subclones were isolated, expanded and preserved for future studies and used as (no-XIST) controls in various contexts (Chiang et al.,2018; Czerminski and Lawrence,2020; Moon and Lawrence,2022). iPSCs were maintained on vitronectin-coated plates with Essential 8 medium (ThermoFisher) and tested periodically for mycoplasma. Cells were passaged every 3-4 days with 0.5mM EDTA. Cell lines were verified for appropriate chromosome 21 number by FISH for a chr21 gene (e.g. *APP)* before each series of differentiations, and trisomy 21 status confirmed by RNA sequencing transcriptomics.

Neural differentiations were performed as previously described (Chambers et al.,2009; Cao et al.,2017) with some modifications. Briefly, iPSCs were dissociated into single cells and plated at a density of 50,000 cells/well in a vitronectin-coated 24-well plate with 10µM of the ROCK inhibitor Y-27632 (Tocris Bioscience). The next day, media was changed to Neural differentiation media (NDM) consisting of 50% DMEM/F12, 50% Neurobasal, 0.5X Glutamax, 1X N-2 supplement, 1X penicillin/streptomycin (all from ThermoFisher), and supplemented with 2uM DMH1 and SB431542 (both from Tocris Bioscience). After 14 days, cells were broken into clumps after EDTA treatment and cultured in suspension for 7 days in NDM. On day 21 or 28 of differentiation, neurospheres were dissociated into single cells with StemPro Accutase (ThermoFisher) and plated onto coverslips (Electron Microscopy Sciences) coated with Matrigel (Corning) at a density of 25,000-50,000 cells/coverslip and fed every 2-3 days with Neuron media consisting of Neurobasal, 1X N-2, 0.5X B-27 without vitamin A, 1X penicillin/streptomycin, 1X Glutamax (ThermoFisher), 0.3% Glucose, 10ng/ml GDNF (Peprotech), 10ng/ml BDNF (Peprotech), 10ng/ml ascorbic acid (Sigma-Aldrich), and 1µM cyclic AMP (Sigma-Aldrich). Doxycycline diluted in distilled water was added to the culture media starting at various time points at a concentration of 500ng/ml. In cultures where NSCs were synchronously differentiated to neurons, compound E (EMD Millipore) was added for 3 days at day 21 of differentiation at a concentration of 200nM.

### Cerebral organoid differentiation

Lancaster protocol: organoids were generated as previously described (Lancaster et al.,2013; Lancaster and Knoblich,2014). Briefly, iPSCs were dissociated into single cells and plated at a density of ~9,000 cells/well in 96-well round-bottom ultra-low attachment plates (Corning) in iPSC media containing 4ng/ml thermostable FGF-2 (Millipore) and 50µM Y-27632 (Tocris Bioscience). After 6 days, organoids were transferred to ultra-low-attachment 24-well plates in N2 and heparin-containing neural induction media. Organoids were embedded in Matrigel droplets on day 11 of differentiation and grown for 4 days before transferring to an orbital shaker set at ~100 RPM.

Paşca protocol: spheroids were generated as previously described (Pasca et al.,2015) with significant alterations. The first steps of the protocol were performed as above, using a re-aggregation strategy. Cells were re-aggregated in 96-well plates in iPSC media containing 20ng/ml thermostable FGF-2 and 50µM Y-27632. The next day, half the media was exchanged with neural differentiation media (NDM) containing 2uM DMH1 (Tocris Bioscience) and SB431542 (Tocris Bioscience). Organoids were fed with this media every day for 14 days. After 14 days, media was changed to neural media containing 20ng/ml FGF-2 and EGF (Peprotech) as described (Pasca et al.,2015) and moved to ultra-low attachment 24-well plates. From this point forward, organoids were grown on an orbital shaker set at ~100 RPM to improve aeration. At day 32, FGF-2 and EGF were replaced with 20ng/ml of BDNF (Peprotech) and NT-3 (Peprotech) for 18 days. At day 50, organoids were fed every other day with neural media without any supplements.

Qian protocol: forebrain organoids were generated as previously described (Qian et al.,2016) with the following modifications: embryoid bodies were formed by dissociated of iPSCs into single cells and re-aggregating in U-bottom 96-well plates (Lancaster and Knoblich,2014). On day 7, aggregates were transferred to ultra-low attachment 6-well plates (Corning) for Matrigel embedding, and on day 14 the plates were moved to an orbital shaker set at ~100rpm.

### Cell fixation, RNA FISH, and immunofluorescence

For iPSC and monolayer neural culture, cell fixation with 4% paraformaldehyde (PFA) was performed as previously described (Byron et al.,2013). Forebrain organoids were fixed for 30min in PFA at room temperature, washed three times with PBS, and cryopreserved in 30% sucrose/PBS at 4°C overnight. Fixed organoids were embedded in O.C.T. compound (Sakura Finetek), frozen in an ispropanol/dry ice slurry, and sectioned at 14µm on a cryotome. Sections were attached to Superfrost Plus slides (Electron Microscopy Sciences) and stored at −20°C until staining. Prior to staining, sections were rehydrated in PBS for 5min, and detergent extracted in 0.5% Triton X-100 (Roche) for 3min.

RNA FISH and IF were performed as previously described (Clemson et al.,1996; Byron et al.,2013). For RNA FISH and combined RNA FISH/IF in iPSCs and monolayer neural culture, detergent extraction was performed prior to fixation. For IF alone, fixation was performed prior to detergent extraction. The XIST probes used were G1A (Addgene plasmid #24690; Clemson et al., 1996) and a Stellaris FISH probe (Biosearch Technologies, SMF-2038-1), which was used according to the manufacturer’s instructions. The APP probe is a BAC from BACPAC resources (RP11-910G8). DNA probes were labelled by nick translation with either biotin-16-dUTP or digoxigenin-11-dUTP (Roche). For simultaneous IF and RNA FISH, cells were immunostained normally with the addition of RNasin Plus (Promega) to the incubation buffer and fixed in 4% PFA prior to RNA FISH. The primary antibodies used in this study are provided in Table 2.1.

The conjugated secondary antibodies used in this study were Alexa Fluor 488, 594, and 647. BrdU staining was performed after RNA FISH and subsequent fixation by incubating coverslips or slides at 80°C in 70% formamide in 2X SSC for 5min (coverslips) or 30min (cryosections on slides) followed by dehydration in 70% and 100% cold ethanol for 5min each and standard IF. Primary antibodies used in these experiments are listed in Table 1.

**Table 1.**
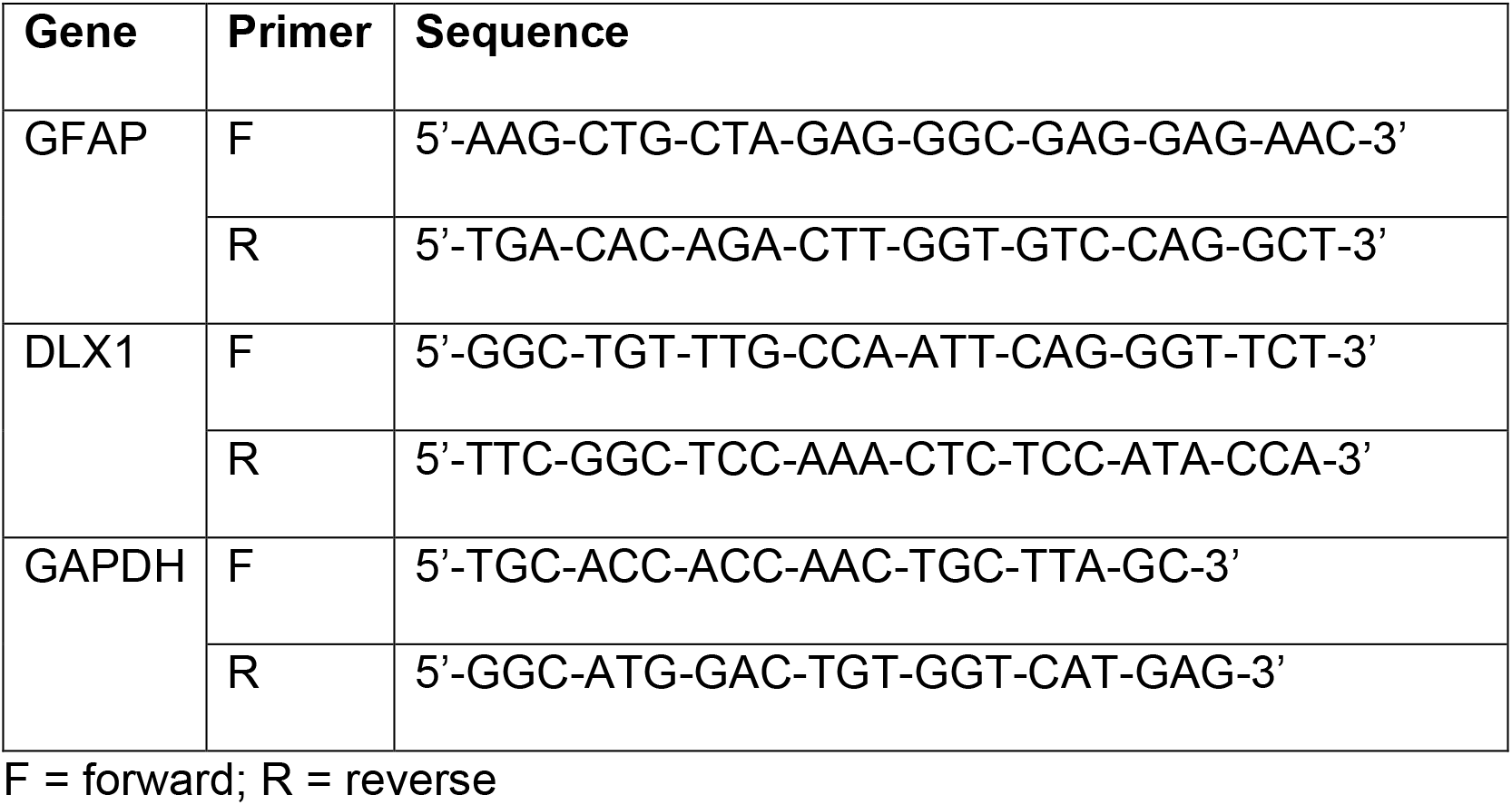
qPCR primers.

### RNA isolation, cDNA library preparation, and high-throughput sequencing

Whole organoids were washed once with 1X PBS and placed into 2ml microcentrifuge tubes containing one 5mm steel bead (Qiagen) and 1ml of Trizol reagent (ThermoFisher). These samples were homogenized using the TissueLyserII instrument (Qiagen) on the P1 setting. Beads were then removed using a magnet and samples were either stored at −80C or RNA extraction, DNAse treatment, and RNA cleanup was performed immediately.

RNA was extracted using TRIzol reagent (ThermoFisher) according to manufacturer’s instructions. RNA samples were cleared of contaminating genomic DNA by DNAse I (Roche) treatment for 1hr at 37°C. RNA cleanup and DNAse I removal was performed using RNeasy MinElute columns (Qiagen) according to manufacturer’s instructions. Clean RNA was assessed for quality on an Advanced Analytical Fragment Analyzer. All samples had an RQN > 7.5 and strand-specific sequencing libraries were prepared using the NEBNext® Ultra™ II Directional RNA Library Prep Kit for Illumina® in conjunction with the NEBNext® Poly(A) mRNA Magnetic Isolation Module and NEBNext® Multiplex Oligos for Illumina® (New England Biolabs).

Sequencing was performed by the UMass Medical School Deep Sequencing Core Facility on the Illumina HiSeq4000 platform to a depth of ~8 million reads/sample in the case of the large organoid experiment or on the NextSeq instrument to a depth of ~30 million reads/sample in the case of the pilot experiment.

### RNA sequencing analysis

Reads were aligned to the hg19 human genome build (GRCh37) using hisat2 (v2.0.5). Reads were counted to genes using the feature Counts function of the subread package (v1.6.2). Within R, the DEseq2 package was used to normalize reads between samples and determine significantly differentially expressed genes. Significance in Figure 3D was determined by performing multiple comparison correction on all expressed chr21 genes (n=125) and setting an FDR of <0.1. The ggplot2 package was used to generate most graphs.

Cell type representation deconvolution was performed using the BSeq-SC algorithm (Baron et al.,2016). Pre-averaged pseudobulk estimates of single cell sequencing data from Quadrato (Quadrato et al.,2017) were used as the basis vectors for deconvolution. The top 20 marker genes ranked by p-value for each cell cluster were used to determine cell type representation estimates.

The quasi-likelihood test in the edgeR package for R was used to determine differential gene expression. Replicate samples and repeated differentiations of the same cell line were summed together to form a 3 vs. 3 comparison. Multiple comparison correction was performed separately for chr21 and non-chr21 genes to determine significant differential expression. For cell type representation correction, the estimated proportion of forebrain cells was included as a covariate in the statistical model.

### Reverse transcription and qPCR

RNA was extracted and processed as described above. Reverse transcription was performed using SuperScript III reverse transcriptase (ThermoFisher) per manufacturer’s instructions and using random hexamers for first strand synthesis. cDNA was then diluted, and qPCR reaction was set up using iTaq Universal SYBR Green Supermix (BioRad) per manufacturer’s instructions with the primers listed in Table 2. The qPCR reaction was performed on the BioRad C1000 Touch thermal cycler. GAPDH was used for normalization and quantification was performed using the ΔΔCt method (Livak and Schmittgen,2001).

**Table 2.** Primary antibodies.

| **Antibody** | **Host** | **Source** | **Identifier** |
| --- | --- | --- | --- |
| NeuN | mouse monoclonal | Millipore | MAB377 |
| Sox2 | rabbit polyclonal | Millipore | AB5603 |
| TUBB3 (Tuj1) | mouse monoclonal | Biolegend | MMS-435P |
| Sox1 | goat polyclonal | R&D Systems | AF3369 |
| GFAP | rabbit polyclonal | Chemicon | AB5804 |
| PAX6 | rabbit polyclonal | Biolegend | 901301 |

### Aβ Analysis

48-hour old media was removed from tissue culture wells containing individual organoids. Media was immediately placed on ice and centrifuged at 2000rcf for 5 minutes to remove cell debris. Media supernatant was stored at −80C. ELISA was performed using the ultrasensitive Amyloid beta ELISA kit from Invitrogen (Paina et al.,2011KHB3544) per manufacturer’s instructions with media samples diluted 1:2 in standard diluent buffer. (The large organoid study determined if Abeta 40 had increased, during a period when the Abeta 42 ELISA kit was unavailable.) Plates were read at 450nm using a BioTek EL800 microplate reader. Graphpad Prism software was used to create a 4-parameter standard curve after subtracting background absorbance. Individual samples were normalized to total RNA extracted from the corresponding organoid harvested on the same day as the media was collected, as detailed in the RNA extraction section.

## Results

The results below detail the progression of experimental design improvements based on results of earlier observations. The last and largest experiment was formulated from lessons learned from our initial cerebral organoid studies, which highlighted the need to address several sources of variation that are often present but not due to trisomy 21, most of which are also relevant to human disease modeling with iPSCs generally. DS studies have reported numerous phenotypic and transcriptional differences attributed to trisomy in human or mouse neural tissues and cells, however it is often difficult to know whether other potential differences between samples have been ruled out. In our initial smaller experiments we found intriguing differences, so we then sought to determine whether this could be accounted for by variation between organoids, experiments, or cell lines, even isogenic cell lines. This led us to focus on quantitative transcriptome analyses using an expanded experimental design of isogenic organoids. While the need for large numbers to control for variation between organoids was not entirely unanticipated, our results indicate that the number of isogenic iPS lines needed to confidently investigate a potentially subtle neurodevelopmental phenotype was more than we initially expected.

### Evaluating three approaches for generating cerebral organoids with DS iPSCs

Several 3D cell culture models of cerebral development have recently been developed, each with its own set of advantages and drawbacks. Most significantly, protocols differ in the usage of exogenous patterning molecules. Lancaster (Lancaster et al.,2013; Lancaster and Knoblich,2014) utilize a protocol with minimal patterning and make use of the “default fate” of differentiating pluripotent cells to become rostral neuroectoderm. This results in the generation of several cerebral cell types, including meninges, choroid plexus, and cortical zones including self-organizing neural stem cell niches and surrounding neurons. The potential advantages of minimal patterning in generating a model of neurodevelopmental disorders include diminishing the potentially overriding effects of non-physiologic levels of patterning and mitotic factors on a subtle defect in differentiation and/or proliferation. Additionally, modeling a wide range of cell types that could be involved in disease pathogenesis in a dense 3D environment could reveal phenotypes absent in monoculture. On the other hand, the broader range of cell types produced may include those that are likely of peripheral relevance to cognitive disability.

Because of this, protocols based on advances in developmental neurobiology and 2D culture techniques have utilized patterning molecules to establish region-specific 3D models of human neurodevelopment. In particular, Paşca (Pasca et al.,2015) utilizes dual-SMAD inhibition, high concentrations of the mitogens FGF2 and EGF, as well as the neurotrophins BDNF and NT3 to generate spheroids that include only cortical-like cells, including both neurons and astroglia. A third protocol utilizes SMAD inhibitors as well as mild WNT signaling activation, which ameliorated apoptotic cell death and potentially further dorsalized the organoids (Qian et al.,2016; Qian et al.,2018).

In order to identify a model of DS neurodevelopment that produced consistent results and was tractable in our hands, we evaluated three protocols using isogenic iPSCs with trisomy 21 and their disomic controls. First, we generated unpatterned cerebral organoids, as previously described (Lancaster et al.,2013; Lancaster and Knoblich,2014). This protocol generated large ventricular-like zones with tightly packed neural stem cells surrounded by outwardly migrating postmitotic neurons (Figure 1A). However, these structures constituted a minority of cells in each organoid, which were largely composed of self-organizing cells in a non-ventricular pattern, reminiscent of choroid plexus-like tissue as well as other cells lacking clear organization (Figure 1A). Because we aim to examine the effects of trisomy on cortical neurogenesis, we decided that this protocol was not sufficiently reproducible to achieve this goal.

**Figure 1.**
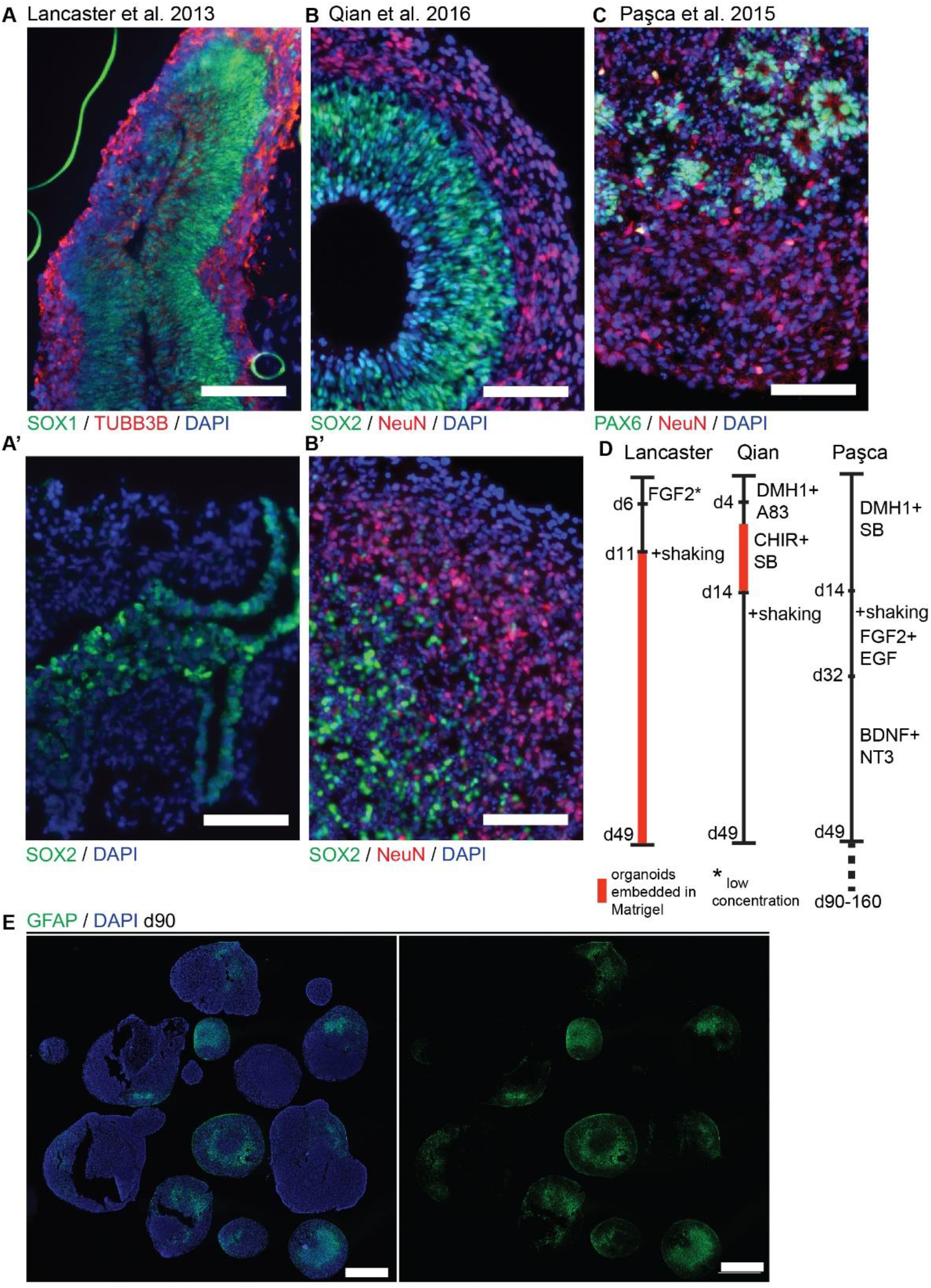
Evaluating three approaches for generating cerebral organoids with DS iPSCs. A-C) Immunofluorescence photomicrographs of representative cortical regions in three selected organoid generation protocols. A’) Non-cortical region in organoid generated with Lancaster protocol, which resembles choroid plexus in organization. B’) Cortical region of organoid generated with Qian protocol but lacking the distinct radial organization seen in (B). D) Visual summary of protocol generation protocols. All protocols utilize the same first step of single-cell dissociation and re-aggregation in 96-well plates. E) Example of 90-day organoids generated with Paşca protocol and demonstrating robust GFAP expression, suggestion formation of astrocytes. Wide variability between individual organoids can be seen, which makes quantification of cell representation difficult. Scale bars are 100μm in (A-C) and 1mm in (E).

Next, we created forebrain organoids with minor modifications (see methods) (Qian et al.,2018). After ~50 days, these organoids formed a large number of large, well-organoids ventricular-like zones (Figure 1B). The organoids were largely comprised of VZs with very few regions that showed organization of a different cerebral cell type. These organoids demonstrated a particularly striking contrast between NSC-containing VZs and surrounding neuron-containing regions. This protocol may be favorable for examination of early VZ formation and the cell dynamics within VZs. However, in our experience some batches generated robust VZ-containing organoids, while others did not produce the characteristic morphology, potentially due to inconsistencies in Matrigel embedding and subsequent disembedding (Figure 1B). This, along with the increased labor required to generate these organoids with Matrigel embedding, motivated us to find another protocol that could generate organoids with higher throughput.

To this end, we generated cortical spheroids, this time with some significant modifications in order to consistently generate organoids in our hands (Pasca et al.,2015). Most significantly, the exposure to SMAD inhibitors was increased from one to two weeks, and the subsequent mitogen and neurotrophin treatments were thus delayed by one week. Nevertheless, this protocol produced large spheroids containing smaller VZ-like zones along with some unorganized progenitor containing areas (Figure 1C). A visual summary of the organoid differentiation protocols tested in this study is provided in Figure 1D. Additionally, prolonged culture generated significant numbers of GFAP-expressing astrocytes (Figure 1E, S1), production of which is limited in other protocols due to the lack of a progenitor expansion step. Together, these favorable characteristics encouraged us to utilize this organoid generation protocol going forward to examine the effects of trisomy on neurodevelopment in aged organoids.

After early attempts to analyze potential differences in cell type representation using histological methods, we quickly came to the conclusion that the still-large degree of variability from organoid to organoid makes accurate quantification a particularly difficult task (Figure 1E). For this reason, we turned to bulk RNA sequencing to deconvolve cell type representations of trisomic and disomic organoids.

### Initial studies identify differences in cell representation between small pools of disomic and trisomic organoids

Our first pilot RNA sequencing experiment in organoids used bulk RNAseq of 10 organoids aged for 160 days, five from an isogenic trisomic (parental) line and five from the euploid control line. Use of isogenic lines avoids differences in genetic background, and the comparison of subclones of the same iPS line avoids differences in the iPS reprogramming process or the somatic cell of origin. The overall strategy was to generate bulk sequence data and use published gene sets for different cell-types to deconvolve the cell-type representation of the bulk sequencing. To evaluate the variation between individual organoids in this first experiment, we sequenced the 10 organoids individually.

We generated RNAseq data for the 10 organoids, each sequenced to a depth of ~30 million reads with 100bp paired-end reads, which provided strong quantification of mRNA levels for each gene, as evidenced by the reproducible difference in chr21 expression between trisomic and disomic organoids (Figure 2A). Further analysis of the transcriptome differences between disomic and trisomic organoids indicated that, in addition to changes in upregulation of many chromosome 21 genes (red), there were widespread changes in expression levels of non-chromosome 21 genes between these samples (Figure 2B). As described in the introduction, widespread differences in non-chromosome 21 gene expression is a common finding in published studies of DS.

**Figure 2.**
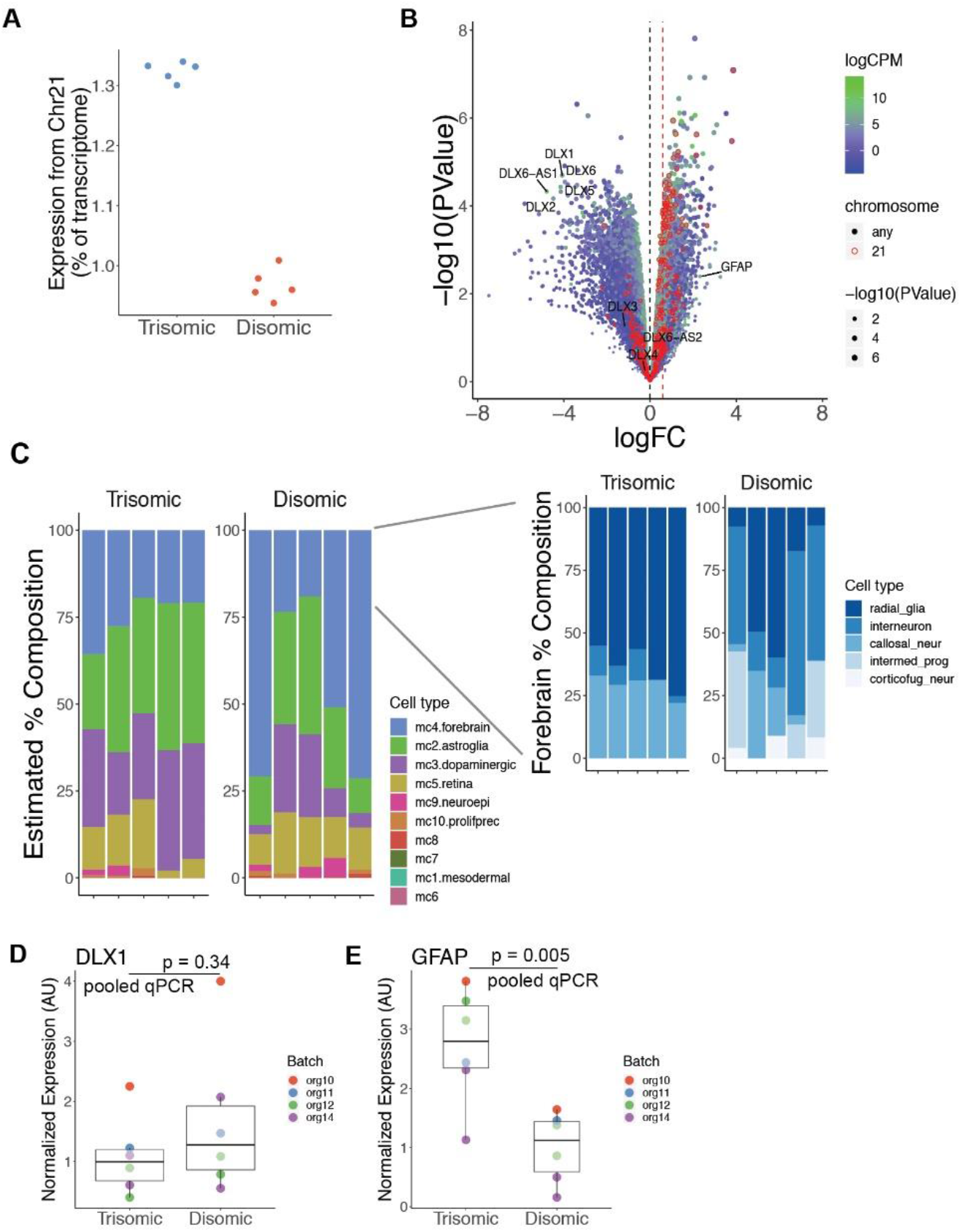
Initial studies identify differences in interneuron or glia cell representation between small pools of disomic and trisomic organoids. A) Fraction of chromosome 21 reads in each individual organoid sequenced. Trisomic organoids have close to the expected 1.5-fold increase in chromosome 21 expression compared to disomic organoids. B) Volcano plot comparing 5 trisomic vs. 5 disomic organoids. Chr21 genes are circled in red and expression level is signified by colors ranging from blue (low expression) to green (high expression). DLX family genes and GFAP are labelled. C) Estimated composition of each organoid into cell types using defined gene sets (further described in methods section). D-E) Pooled qPCR quantification of DLX1 (D) and GFAP (E) expression, normalized to GAPDH. Each dot represents a pool of ~8 organoids. Colors represent independent organoid differentiations.

Most notably, analysis of differentially expressed non-chr21 genes between the two conditions showed many DLX family genes, each identified as downregulated on average in the trisomic organoids (Figure 2B). These genes are well-known for their involvement in the specification and migration of ventral forebrain-derived interneurons (Anderson et al.,1997; Stuhmer et al.,2002; Cobos et al.,2007; Paina et al.,2011), and there have been mixed reports in human samples and cell models of whether interneuron generation is decreased or increased due to trisomy (Ross et al.,1984; Bhattacharyya et al.,2009; Huo et al.,2018; Xu et al.,2019).

To further investigate whether other interneuron-related genes followed the same pattern, and to determine whether other cell types also had altered representations in trisomic organoids, we utilized marker gene lists recently generated by single-cell RNA sequencing of human cerebral organoids (Quadrato et al.,2017). There could be a statistically significant difference on average between these two samples of five versus five organoids, but that does not itself address whether this reflects a consistent difference in neurodevelopment between these trisomic versus control cells. Deconvolution of cell type representations for each individual organoid demonstrated overrepresentation of forebrain-derived cells in the disomic condition, largely driven by an increase in interneuron generation (Figure 3C). Importantly, this effect was driven by three disomic organoids which had large numbers of this cell type, whereas two disomic organoids had similar interneuron composition to the trisomic organoids. A second difference in cell-type representation was observed for radial glia cells, which were overrepresented in trisomic organoids, with a corresponding increase in GFAP expression, a gene also expressed in astrocytes, in these samples (Figure 2B). Thus, despite using a directed organoid generation protocol, significant variability between individual organoids in interneuron formation weakens any conclusions that can be drawn on this point.

**Figure 3.**
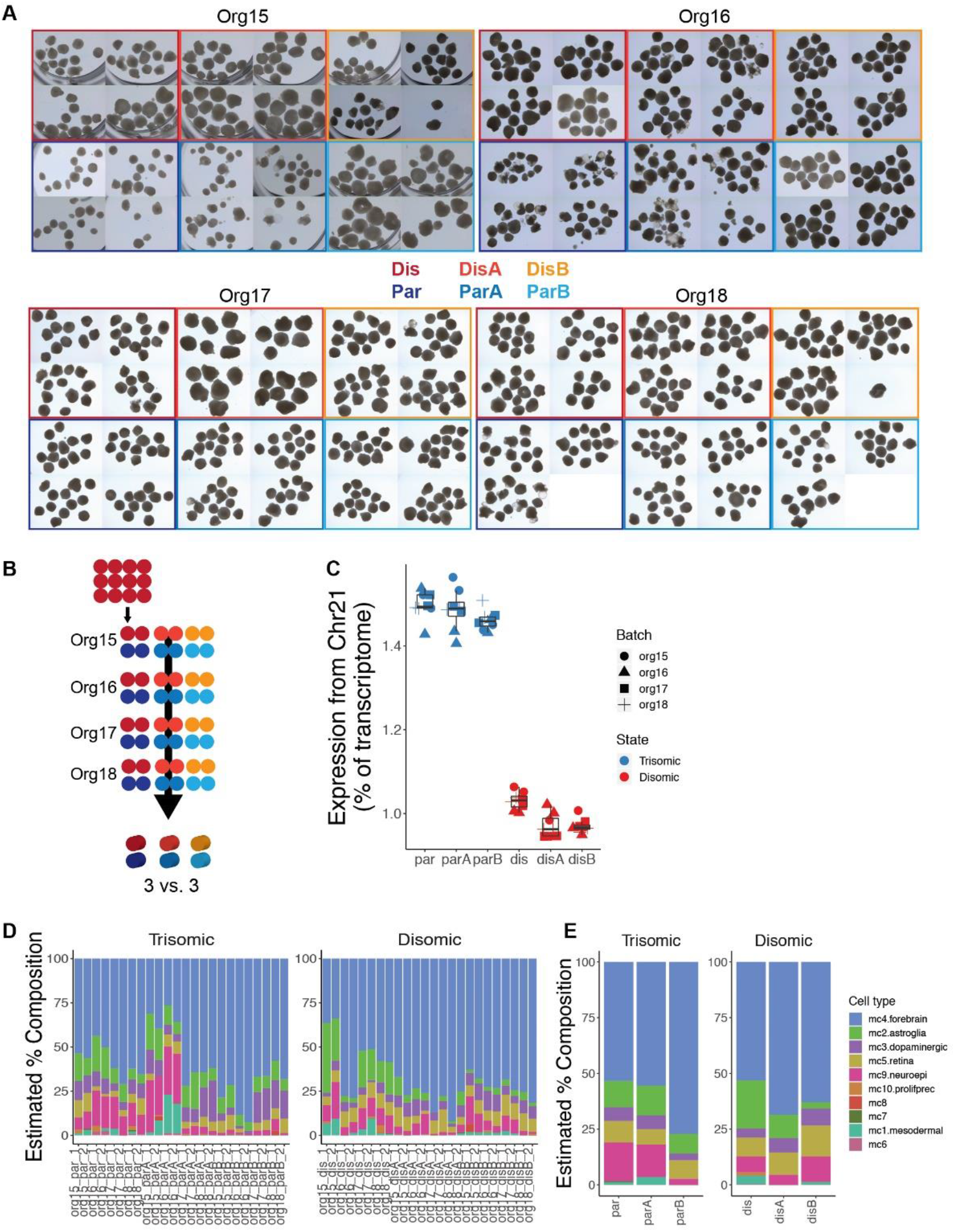
Expanded experimental design to discriminate differences due to trisomy 21. A) Micrographs of nearly all organoids generated in this experiment. Independent differentiations are signified by “org”, isogenic trisomic lines by “par”, and isogenic disomic lines by “dis”. B) Schematic of samples generated. Each of the 48 dots represents 12 organoids and one sample for sequencing, while the 3D cylinders signify *in silico* collapsing for statistical comparison. C) Fraction of chromosome 21 reads for each pooled sample sequenced. D) Estimated composition of each sample into cell types using defined gene sets. E) Estimated cell-type composition of collapsed samples used for statistical comparison.

In an attempt to minimize differences between individual organoids, and to investigate whether similar findings could be identified in younger organoids, we generated several batches of organoids grown for 90 days and examined them in pools of ~8 organoids by RT-qPCR for a prominent marker of interneurons, DLX1, as well as for GFAP. We found that DLX1 expression remained highly variable between experiments and inconsistently showed an increase in disomic organoids over trisomic, although we note that higher expression in organoids from the parental (trisomic) line was not seen in these experiments (Figure 2D). On the other hand, GFAP expression was consistently upregulated in the trisomic organoids, by about 3-fold (Figure 2E). This indicates that either the trisomic cells generate more radial glia and/or astrocytes, or that these cell types produce higher levels of GFAP mRNA in the trisomic condition.

### Expanded experimental design to discriminate differences due to trisomy 21

The above findings provide some evidence of differences in organoid development that correlates with trisomy, but questions remain as to whether these differences could reflect other sources of variability. Therefore, we greatly expanded the experimental design (Table 3) in order to increase the power to discriminate differences due to trisomy from differences due to variability between individual organoids, different experiments, or even between isogenic cell lines. In this experiment we generated a total of over 1,100 organoids (Table 3, Fig. 3A) from three trisomic and three disomic isogenic iPSC lines, which were derived from the same iPS cell reprogramming event. To minimize effects of individual organoid differences, we examined pools of 12 organoids, four pools per cell line, and repeated this scheme in four independent batches of organoids

**Table 3.**
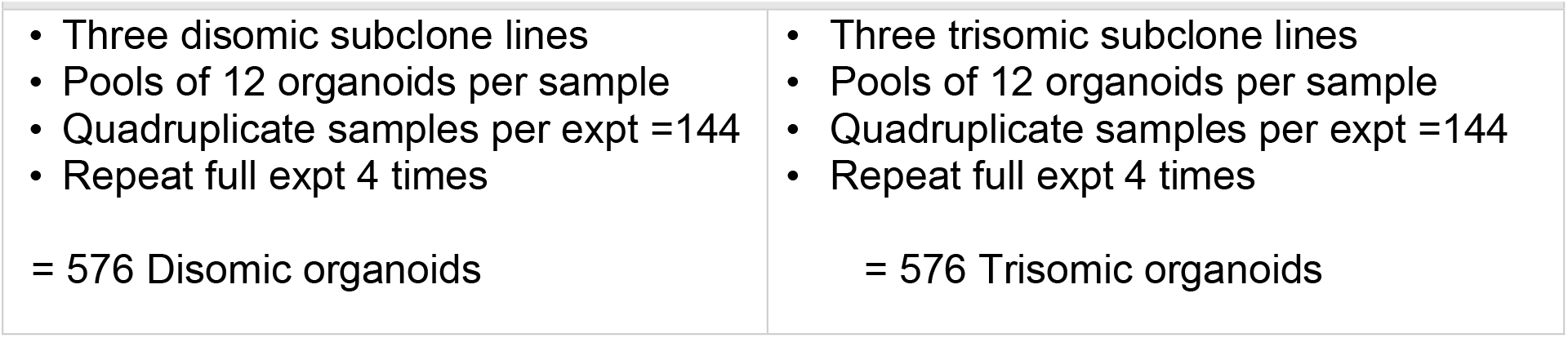
Expanded organoid experiment. Minimize sources of variation >1100 isogenic organoids

Roughly half of these organoids were used for bulk RNA sequencing, with 2 pooled samples per cell line per batch, for a total of 48 samples (Figure 3B). The remaining organoids were frozen for histology and media preserved to assay for Aβ secretion, and for other future analyses on parallel samples.

Initial sequencing analysis confirmed expected differences in chromosome 21 expression, with all trisomic lines having ~1.5-fold higher levels of chromosome 21 transcripts compared to disomic lines (Figure 3C). We next set out to use this transcriptome data to determine the cell type composition of each sample using the same list of gene markers described above, based on published single-cell seq studies defining specific gene sets. We found that nearly every sample was mostly composed of forebrain-type cells, with significant contributions from astroglia, dopaminergic neurons, neuroepithelial cells. Surprisingly, this analysis revealed that some organoid samples contained a subset of mesoderm-derived cells (Figure 3D), suggesting some degree of off-target differentiation.

Notably, there was no consistent or statistically significant difference in the proportions of the categorized cell-types between the disomic and trisomic states (Figure 3E). Differences in cell-type representations between cell lines of the same state (disomic or trisomic) were detected, but these were not consistent between the disomic versus trisomic lines. This cell type variability can also be visualized by IF staining for the astrocyte marker GFAP in organoids from the 6 isogenic lines (Figure S1), which shows significant inter and intra cell line variability. Comparison of results for a given line between independent differentiations suggested that some variation appears sporadic but may also reflect inherent epigenetic differences between even isogenic cell lines which may evolve in culture (see Table 2, Discussion). We cannot rule out that the variability detected between cell lines may mask the presence of more subtle differences in the propensity of disomic and trisomic organoids to form different neural cell types.

Since the presence of the extra chromosome may confer increased cell stress (Oromendia et al.,2012; Sheltzer et al.,2012; Bonney et al.,2015) or cell senescence (Nawa et al.,2019; Meharena et al.,2022) that could reduce general cell proliferation, we also briefly considered if there was a difference in the overall size of the trisomic vs disomic organoids, as found in a recent study (Tang et al.,2021). The numerous pools of 90-day organoids were photographed (as illustrated in Figure S2.A) and measured, and results summarized in Figure S2.B (Supplement). Based on results of all trisomic and all disomic organoids, the average size is smaller for the trisomic lines. However, as indicated in the graph showing results for each of the six cell lines, there was variability between lines (and experiments) that did not cleanly correlate with T21 status. Hence, more isogenic trisomic and disomic lines would need to be studied to determine if the average reflects a generalizable effect. (Note that if just two disomic and two trisomic clones were compared, results would depend on which cell lines/experiments were compared.) Our own observations comparing normal and DS human fibroblast lines suggested that trisomic cells were more susceptible to replicative senescence (Swanson,2014) (Dissertation Chapter IV), consistent with other recent evidence that trisomy 21 can increase cell senescence under stress (Oromendia et al.,2012; Marcovecchio et al.,2021). However, while senescence was not directly studied in this stem-cell based study, we did not detect increased expression of p16 or p21, markers of senescent cells.

A theme of recent transcriptome studies in DS cells and tissues is the finding of extensive transcriptome-wide differences between trisomic and euploid samples. This is also suggested in our initial experiment comparing individual organoids from trisomic and disomic iPSCs. To determine whether this expanded organoid experiment would affirm a similar finding, we examined differentially expressed genes (DEGs) between disomic and trisomic organoids. An important aspect of this statistical analysis was to avoid amplification of clone-specific differences by treating repeated measurements from different experiments on the same cell sample as four independent measurements, a form of pseudo-replication which results in inappropriately inflated p-values. Thus, our analysis collapsed replicate samples and samples from different differentiations of the same cell line and compared organoids generated from the 3 trisomic lines to 3 isogenic disomic lines (Figure 3B).

This analysis detected strong upregulation of genes across chromosome 21, with 159 of 250 expressed chr21 genes meeting statistical criteria for differential expression, generally at or near the 1.5-fold level expected in trisomic cells (Figure 4A). Interestingly, a notable exception was the RWDD2B gene, which was over 13-fold upregulated in trisomic samples. The most striking finding, however, was that despite the robust detection of differentially expressed chromosome 21 genes, no non-chr21 transcriptome changes were detected (FDR<0.1) in this greatly expanded experiment. This suggests that the genome-wide differences found in the smaller-scale experiment above (Fig 2B) may reflect biological differences between the cell lines compared, rather than due to trisomy. However, two astrocyte marker genes, GFAP and AQP4, both showed a trend towards upregulation in trisomic samples, although these did not meet this significance threshold, likely due to variable expression between samples (Figure 4C). Hence, these results do not contradict other studies that have reported increased astrocytes, since inter-cell line variation may have muted the significance of a difference due to T21, as is hinted in our results.

**Figure 4.**
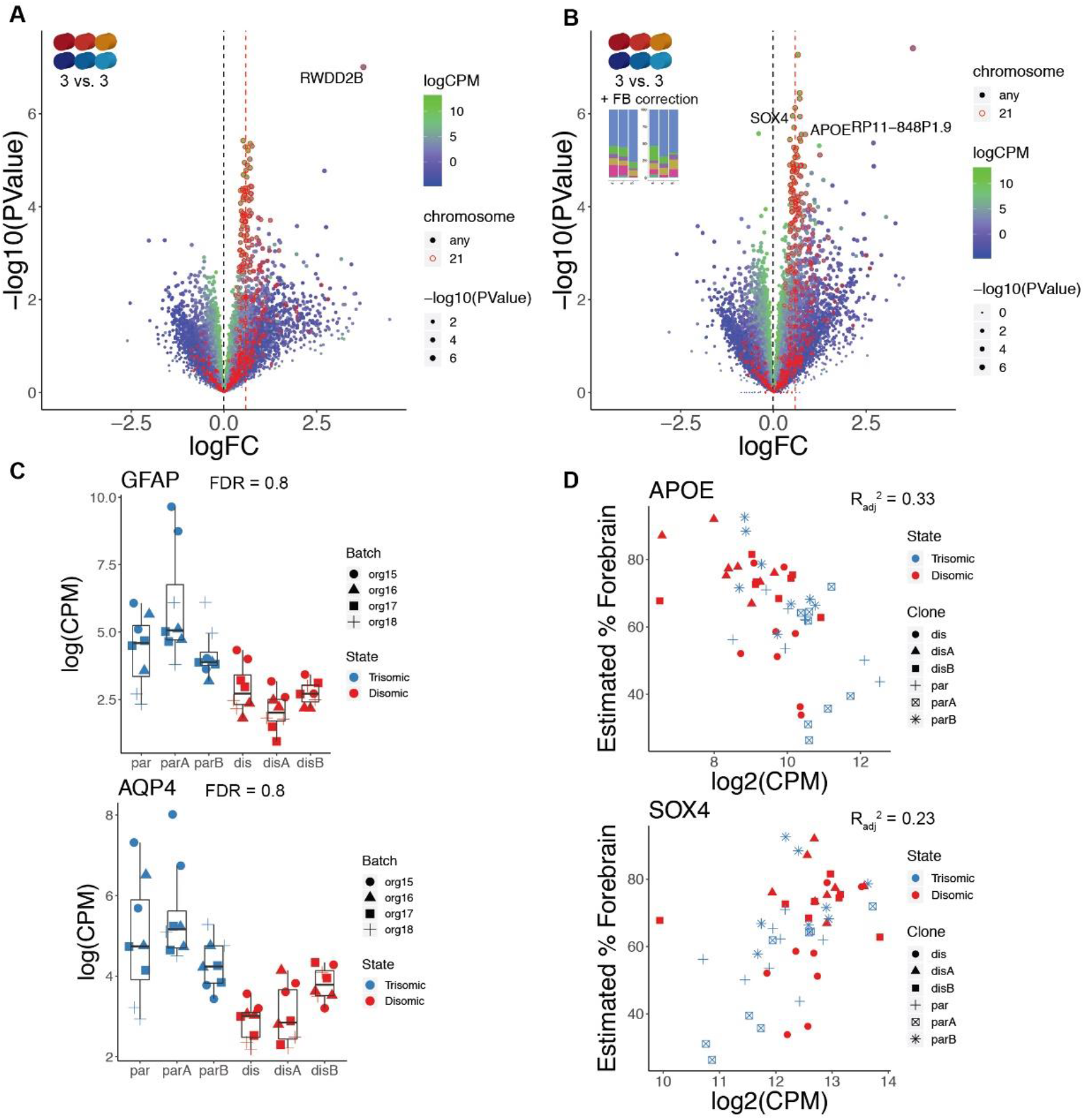
Genome-wide transcriptome analysis of expanded organoid experiment. A) Volcano plot of collapsed 3 vs. 3 comparisons of trisomic and disomic conditions. Chr21 genes are circled in red. After FDR correction for the number of chr21 vs non-chr21 genes, B) Volcano plot of the same comparison as in (A) but including the degree of forebrain representation as a covariate for each sample. The 3 non-chr21 genes that are identified as statistically significant (FDR<0.1) are labelled. C) Individual gene plots for two astrocyte marker genes, GFAP and AQP4 which do not meet statistical difference thresholds but trend towards higher levels in trisomic samples. D) Expression level in each sample of two of the non-chr21 genes identified in (B) as statistically significant plotted against estimated forebrain composition demonstrating strong correlation.

The paucity of non-chr21 DEGs detected in this expanded study fits with the lack of consistent differences in cell type representation shown above, since consistent differences in cell-type representation would likely result in differential expression of many cell-type specific genes. On the other hand, inconsistent variation in the cell type representations between all the cell samples (not correlated with trisomy/disomy) might increase gene expression variability that potentially could obscure any subtle changes in gene expression between the two groups. In an attempt to correct for this possible effect, we normalized the data in each sample for the dominant cell type, the proportion of forebrain cells. Importantly, this correction increased the number of significantly DE chr21 genes to 175 (Figure 4B), indicating an increase in power due to decreased variance by correcting for differences in cell type representations between cell lines. Off of chr21, three hits emerged, SOX4, APOE, and a pseudogene (RP11-848P1.9) that was expressed at low levels. As expected for the emergence of significant genes when correcting for cell type representation, SOX4 and APOE are correlated, positively and negatively, respectively, with the estimated proportion of forebrain cells (Figure.4D). Of these, SOX4 expression was downregulated in trisomic samples, whereas APOE and RP11-848.9 were more highly expressed. While changes in these genes were only detected after computational correction for forebrain representation, they suggest possible involvement of two potentially important genes that to our knowledge have not been previously implicated in relation to DS (see Discussion).

### Over-production of Aβ is already evident in trisomic organoids

An in-depth study of AD-related cell phenotypes is the subject of a separate study, however we include here limited analysis of *A*β to contrast detection of this neurodegenerative pathology to the neurodevelopmental results. In the large organoid study we isolated media from each organoid pool and analyzed whether an increase in secreted Aβ secreted would be detected in these fetal stage organoids, using ELISA for Aβ 40 (see Methods).. Figure 5 shows data for four experimental replicates collapsed to show results for three trisomic versus three disomic lines (all isogenic). Note that there is variability between isogenic trisomic lines, and between the disomic lines, and some variation between experimental batches; importantly, however the strength of the Aβ phenotype is stronger than the sample variations. Remarkably, the strongest phenotype seen in this study of DS fetal dorsal forebrain development relates to neurodegeneration (Alzheimer Disease) rather than neurodevelopment. In this context it is especially intriguing that one of the two non-chr21 genes found to be differentially expressed in this study is APOE, which is involved in the pathology of AD (Yamazaki et al.,2019).

**Figure 5.**
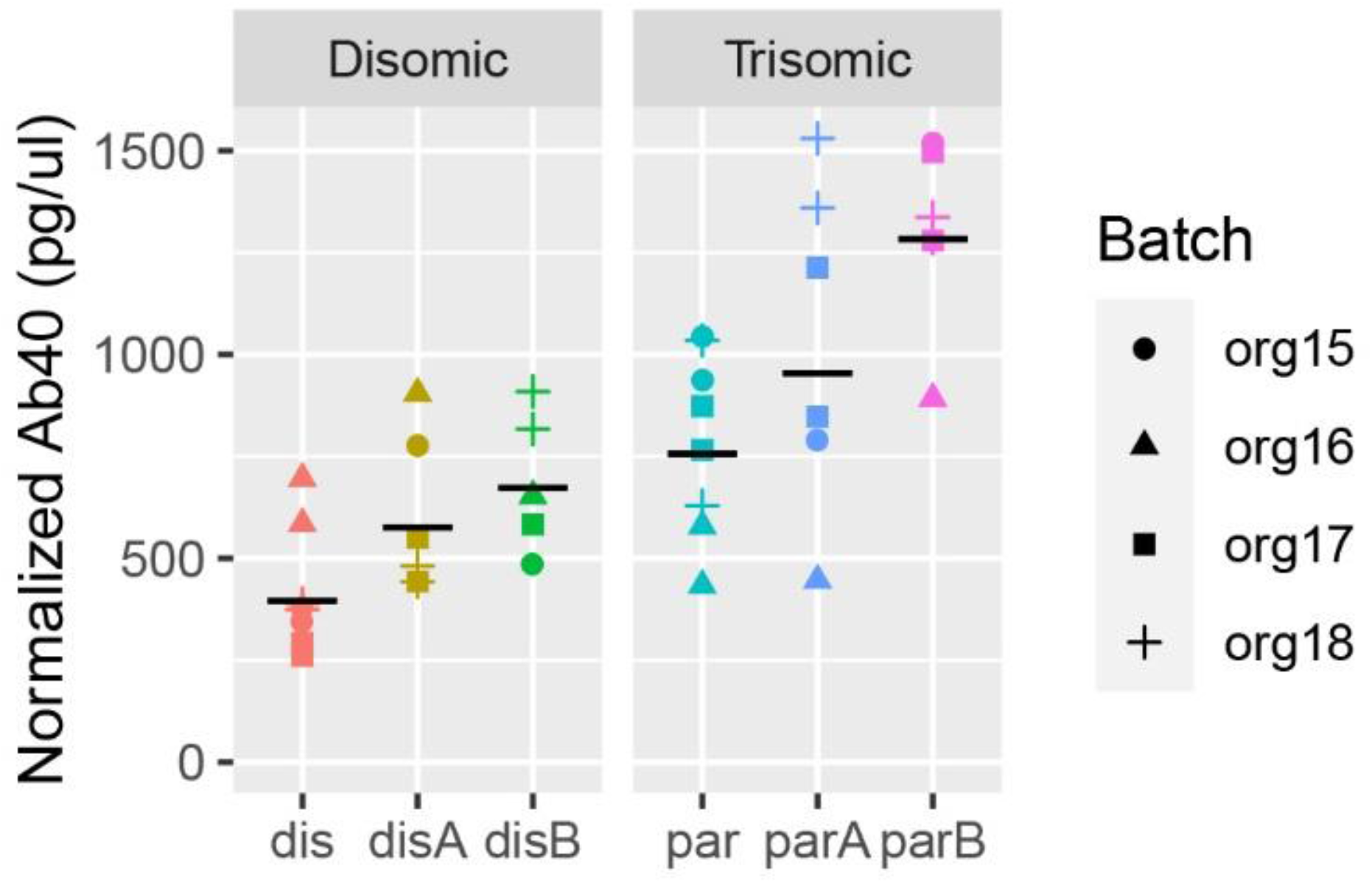
Aβ40 secretion is increased at least 1.5-fold in trisomic organoids. Aβ40 levels were measured in disomic (left) and trisomic (right) organoid media pooled by line and organoid batch and normalized by RNA content. Two media replicate tests were done for each condition. Black line = geometric mean

## Discussion

The overall goal of this study was to use recently developed cerebral organoid technology to shed light on the molecular and cellular pathways altered in early DS neurodevelopment. Understanding how and when brain development and/or function is impacted in DS is critical to assess therapeutic prospects to mitigate cognitive or neurological deficits due to trisomy 21. This study contributes to understanding the extent to which the impact of trisomy 21 on fetal brain development is manifest, as modeled in these 90-day human forebrain organoids. In addition, this study contributes significant technological insights into use of iPSCs and organoids for disease modeling, important to interpretation of results.

Numerous studies report a variety of differences thought due to reflect the impact of trisomy 21 (or its orthologs) in mouse models or human DS tissues and cells. Inconsistencies in findings between studies may be due to differences in the systems examined, or limitations in sample size, etc. Human iPSC studies potentially provide tightly-controlled comparisons between trisomic and euploid cells/organoids in comparable developmental/functional state. However, iPSC cultures are prone to environmental changes that can affect experimental results (Klein et al.,2022) and organoids can exhibit not only individual organoid but batch to batch variability in cell composition (Hernandez et al.,2022). Hence controlling for technical/biological variation in the laborious culture and differentiation of stem cells/organoids is a challenge..Moreover, comparisons can also be impacted by common genetic and epigenetic changes that may evolve during the separate culture of isogenic human pluripotent cell clones and sub-clones [for example, (Hall et al.,2008; Lund et al.,2012; Halliwell et al.,2020)]. Table 4 summarizes several sources of variation that may complicate interpretation of iPSC disease modeling, and summarizes the strategies used in this study to minimize sources of variation.

**Table 4.** Potential sources of variability in iPSC disease modeling.

| **Sources of variability** | **Strategies used in this study to lessen variability** |
| --- | --- |
| Genetic differences between individuals | All Isogenic cell lines |
| Differences in Reprogramming Between isogenic clones from different reprogramming events, from different cells | Subclones from same reprogramming event |
| Differences between “identical” sub-clones<br>B. evolution during culture<br>- Genetic drift<br>- Epigenetic drift<br>B. Freeze/thaw bottleneck<br>-Genetic drift<br>-Epigenetic drift | Expand lines to six (three disomic/three trisomic) |
| Differences between individual organoids | Pooling large numbers of organoids |
| Differences between differentiations | Four repetitions using a semi-directed (relatively consistent) protocol |

As this study progressed from smaller to much larger scale experiments, we worked to minimize or account for these sources of variation on several levels. From the start we used a totally isogenic system, using cell lines that were derived from a single-reprogramming event and cell of origin, since it is known that isogenic iPSC clones from different programming events can show differences in neural differentiation potential (Koyanagi-Aoi et al.,2013). Individual organoids of the same line can be highly variable, so we generated organoids with a semi-directed (less variable) forebrain protocol, and used multiple large pools of organoids per sample in repeated experiments, which we believe adequately controlled for organoid variability. Large numbers of pooled organoids were generated from each of six isogenic lines (three trisomic and three disomic) that are subclones representing the same iPSC reprogramming event. Finally, the entire large organoid production scheme was repeated four times, allowing us to assess and account for variability between experiments.

Due to these efforts, we were able to improve detection of what is a relatively subtle 1.5-fold expected increase in expression for individual chromosome 21 genes, with over 70% of expressed chr21 genes meeting statistical criteria for differential expression due to trisomy. This is substantially more chr21 DEGs than detected in our smaller experiment or in most studies of DS tissues or cells, which, paradoxically, report many more off-chr21 DEGs than found here (Vilardell et al.,2011; Weick et al.,2013; Letourneau et al.,2014; Olmos-Serrano et al.,2016; Mowery et al.,2018). Despite especially robust detection of chr21 DEGs, our largest experiment did not find significant evidence for genome-wide transcriptional deregulation of non-chr21 genes. In our smaller experiment (comparing five organoids each from a trisomic and a disomic line), more genome-wide expression differences were detected between these samples, but, importantly, the larger analysis leads us to reinterpret these results as likely reflecting other sources of biological variability that is real, but we cannot conclude is “dysregulation” due to trisomy 21.

Consistent with a lack of abundant non-chr21 DEGs, the more powerful experimental design did not detect statistically significant differences in cell-type representations between trisomic and disomic organoids. Instead, despite pooling large numbers of organoids to eliminate individual organoid variability, we saw considerable variability between cell lines of the same chr21 state. There was little variability between duplicate samples (pools of 12) for each differentiation experiment (Figure 3-D), and most lines generated similar proportions of cell types in the four differentiation repeats, although one trisomic and one disomic showed more variability between differentiation experiments. Most importantly, the average of all eight pooled samples for each of the six lines (Figure 3-E) showed that differences in cell-type proportions did not correlate with trisomy 21 status. Differences between cell lines (with the same trisomy 21 status) could obscure any subtle changes in cell type representation caused by trisomy 21. In addition, differences between lines could potentially also be conflated with differences due to trisomy.

Organoid technology affords the opportunity to examine development of a greater complexity of cell-types, however that complexity also can introduce analytical challenges. The overall variation in cell-type proportions between samples could weaken the power to identify genome-wide expression differences genuinely due to trisomy 21. We note, however, that chr21 gene changes were strongly detected despite the complexity of organoid samples, indicating that there may indeed be relatively limited effects on the overall transcriptome at this stage. In an effort to minimize noise due to cell-type proportions between samples, we normalized results for the representation of forebrain cells in each sample. Following this correction, just three statistically significant non-chr21 genes were detected. Remarkably, two of these, Sox4 and APOE link to important cellular processes in neurodevelopment and neurodegeneration (AD), respectively, in DS. SOX4 was downregulated in trisomy, whereas APOE RNA increased. SOX4 is an important regulator of neurodevelopment (Kavyanifar et al.,2018), and has been shown to affect oligodendrocyte development by induction of a Notch target gene (Braccioli et al.,2018). This connection to the Notch pathway was especially interesting given our previous finding (Czerminski and Lawrence,2020), using our chromosome silencing system in monolayer neuronal cultures, indicated a strong upregulation of the Notch pathway. Furthermore, missense mutations in SOX4 have recently been identified in patients with intellectual disability and facial dysmorphism (Zawerton et al.,2019) while biallelic deletion has been associated with developmental delay, hypotonia and intellectual disability (Ghaffar et al.,2021), phenotypes also found in DS. The role that SOX4 plays in DS neurodevelopment has not been examined but results here suggest a new hypothesis that, if can be corroborated in other studies, would provide opportunity to explore both the upstream cause (stemming from trisomy 21) of SOX4 dysregulation and its downstream consequences.

Similarly, it seems striking that among just three genes found to differ in trisomic samples was the APOE gene, which, through its isoforms, has been strongly linked to the risk of developing AD (Poirier et al.,1993). While the APOE4 allele confers increased risk of AD and the APOE2 may lower risk, the trisomic (and disomic) cells studied here carry just APOE3 alleles, and thus this would not explain differences in APOE mRNA levels. Interestingly, it has been suggested that the expression level of APOE, which we found elevated in trisomic samples may influence AD pathogenesis independent of isoforms (Huang et al.,2017). Astroglia cells express higher APOE, although preliminary assessmentdid not find the APOE expression pattern to correlate with estimated astroglia proportions in different samples. Hence, the potential dysregulation of APOE, as well as Sox4, require and are being further investigated in specific cell-types, with and without inducible trisomy 21 silencing. While AD-related pathology is not studied in detail here, it is noteworthy that the increased Aβ se cretion is evident in these fetal-stage organoids, despite some variation between lines and experiments. Thus, this neurodegeneration-linked phenotype is less subtle than neurodevelopmental differences in these fetal stage organoids.

Previous studies in iPS-derived DS cells have described a range of findings with many reporting no difference in the neuronal differentiation capacity of DS cells (Shi et al.,2012; Briggs et al.,2013; Lu et al.,2013; Weick et al.,2013; Gonzales et al.,2018). Other studies using unrelated disomic and trisomic iPSCs in a monolayer culture system have demonstrated an increase in the proportion of astroglia formed by trisomic cells (Chen et al.,2014). Another very recent study generated patterned ventral forebrain organoids using DS cells and found an increase in the propensity of trisomic cells to form interneurons, which was correctible by knockdown of a chr21 gene, OLIG2 (Xu et al.,2019). This finding contrasts with previous studies in iPSCs and primary human cells that describe the opposite finding (Ross et al.,1984; Bhattacharyya et al.,2009; Huo et al.,2018). Our findings are not directly comparable to these studies due to significant differences in the organoid generation protocols. We used organoids patterned towards a specific forebrain subregion as in Xu (Xu et al.,2019) in order to decrease variability between organoids and the number of different cell types formed. While this may allow for enhanced detection of disease-specific differences, it could miss differences in specific cell-types not well-represented, such as oligodendrocytes or glial cells. Thus our finding do not contradict these studies, but highlight the challenge that neurodevelopmental differences seen may be impacted by experimental design.

Overall, these results raise caution about false-positive results, but also potential false negative results, arising due to variability from organoid to organoid, batch to batch, and cell line to cell line. Neurodevelopmental phenotypes may be quite subtle and difficult to model with iPSCs, as discussed elsewhere (Soldner and Jaenisch,2012). In the current study we believe we sufficiently minimized the variability between organoids by studying over 500 trisomic and 500 disomic organoids. However, our analysis reveals that in the end, using organoids derived from three trisomic and three trisomic isogenic cell lines and addressing other sources of variability, differences between those cell lines still exist and can over-shadow milder phenotypes. In fact, epigenetic drift in pluripotent cells was earlier demonstrated in our lab and others by demonstrating variability in X-chromatin modifications and *XIST* RNA status in female ES cell lines (Hall et al.,2008; Silva et al.,2008), which we showed evolve even between colonies within the same culture dish. Thus, even isogenic subclones can develop epigenetic differences over time that complicate disease modeling. Increasing the number of isogenic iPSC lines studied from six to perhaps 10-12 might overcome or mitigate this variability, however this is challenging given the need for very large numbers of organoids and repeated experiments as well. For this reason, we have worked to overcome technical difficulties to using dox-inducible trisomy 21 silencing (with XIST) in organoids, and are working to examine the impact of trisomy 21 expression in organoids of the same cell line, which may further illuminate what differences can be directly attributed to trisomy 21.

## Acknowledgements

We thank members of the Lawrence lab for thoughtful discussion, critical analysis, and helpful comments on the manuscript. We appreciate the support of NIH – R35GM122597, R01HD091357, and R01HD094788 to J.B.L.; F30HD086975 and T32GM107000 to J.T.C. J.B.L. also appreciates prior supplemental support from The John Merck Fund.

