## Supplement for "Modeling Down syndrome neurodevelopment with isogenic cerebral organoids"

**Supplementary Material**

**Figure S1.**

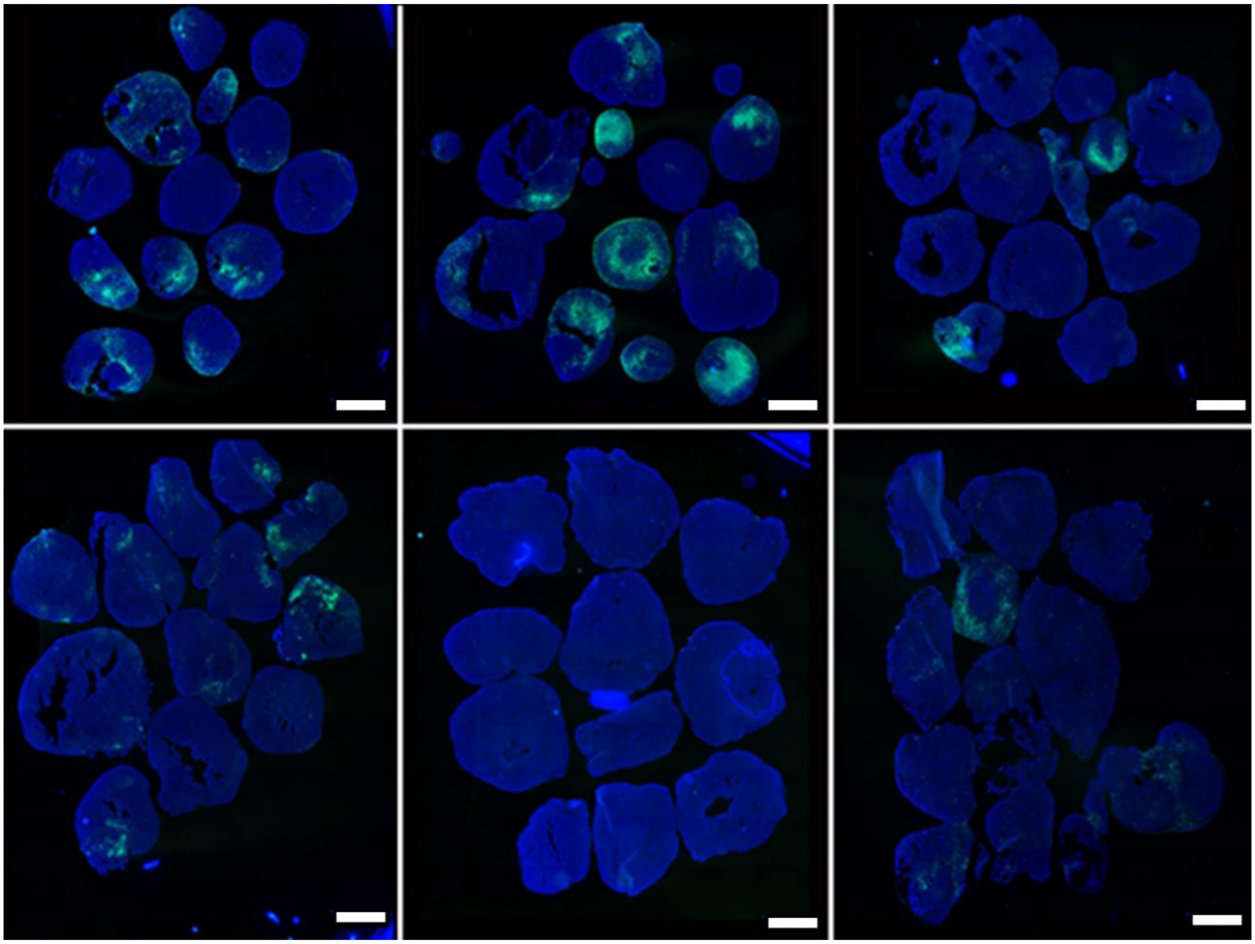

**Fig S1.** GFAP staining (green) of 90-day organoids from each of the 6 cell lines (3 disomic, 3 trisomic) shows variability between lines as well as variability between individual organoids from the same line. Scale bars = 1mm.

**Figure S2**

**A.**

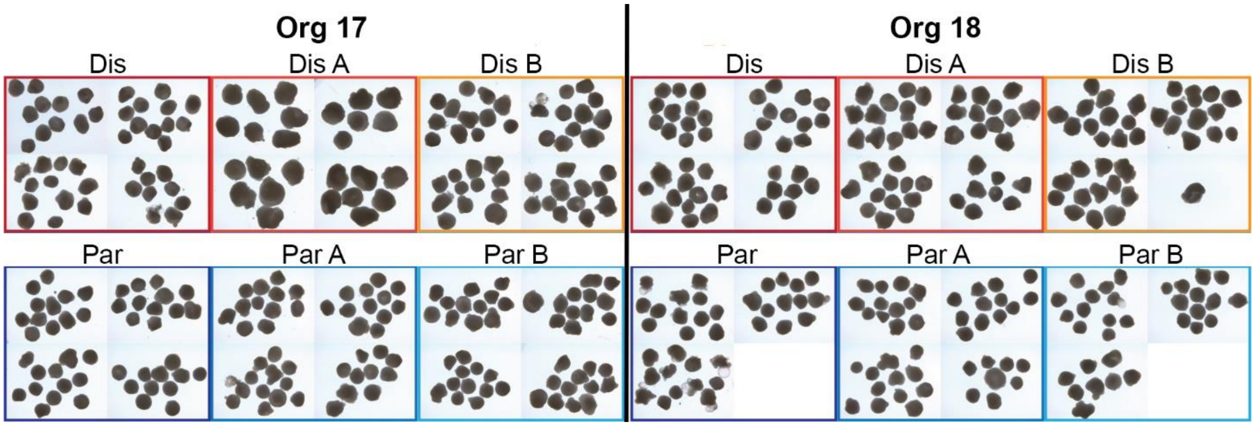


**B.**

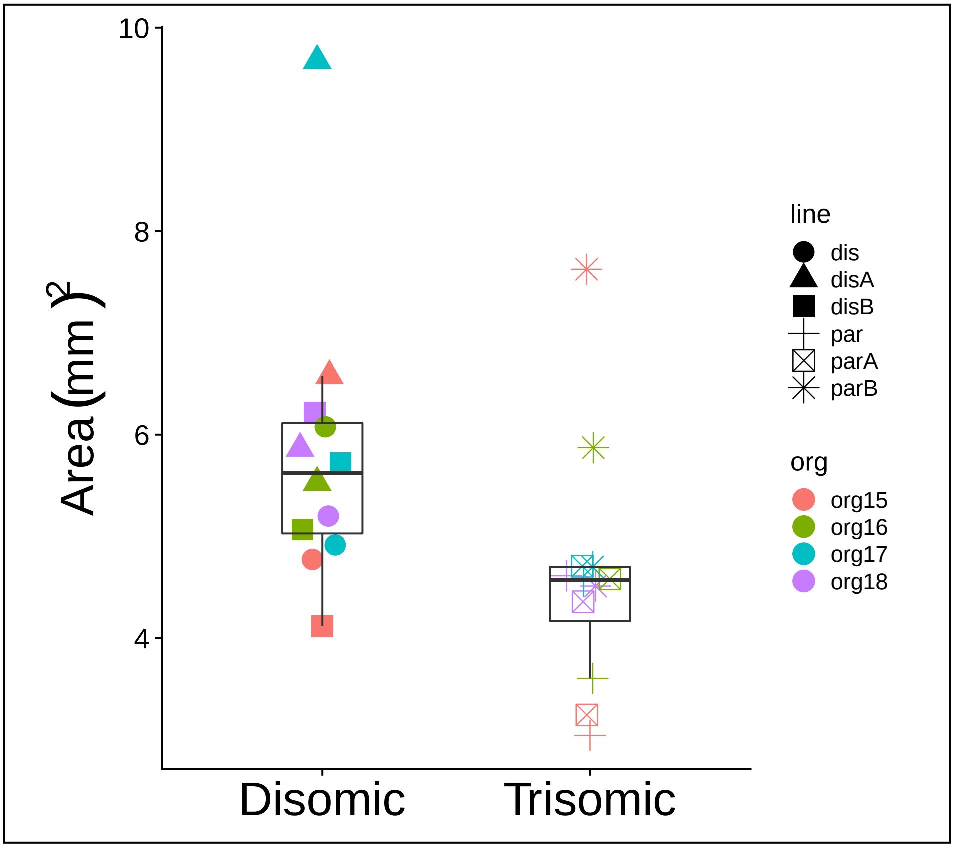


**Fig S2. A.** Examples of 90 day disomic (top panel) and trisomic (bottom panel) organoids from 2 experiments. **B.** Measurements of 90 day organoids from 3 disomic and 3 trisomic subclone lines (4 experiments of each). Data suggest that trisomic organoids may be smaller, but results are inconclusive due to variability between experiments.
